# Rapid and divergent changes in the continental-scale organisation of a short-lived songbird’s migratory strategy

**DOI:** 10.64898/2026.06.16.732301

**Authors:** Joe Wynn, Monika Broniszewska, Alice Edney, Tania Garrido Garduno, Joe Moford, Michał Polakowski, Robert E Rollins, Pablo Salmon, Oscar Vedder, Miriam Liedvogel

## Abstract

It is hard to predict how rapidly songbird migration will change in the Anthropocene. Indeed, since songbird migration is thought to have a strong heritable component, the continental-scale organisation of migratory movement might be seen as fairly inflexible. Perhaps one of the best models for the ecology and evolution of migration is the Eurasian blackcap (*Sylvia atricapilla*) which, as part of a continent-wide effort to characterise blackcap migratory phenotype, we geolocator-tracked from breeding sites in eastern Poland. Rather than migrating in the expected south-easterly migratory direction, these birds migrated south and south-west – suggesting that blackcaps in the east of their range have switched migratory direction. We sought to investigate the extent of this phenomenon using almost a century of ringing data, which confirmed that blackcaps breeding across the entirety of Eastern Europe have indeed almost completely stopped using their historic eastern flyway. Instead, a shorter-distance west-migrating phenotype has emerged, which we find is consistent with warmer winter temperatures opening up wintering sites at more northerly latitudes in the west. We discuss what drives changes in migratory behaviour over short timescales; and consider what this tells us about how migratory information is inherited.

## Introduction

Migration is perhaps the most remarkable adaptation to seasonal changes in the environment, allowing labile animals to escape inclement conditions as the seasons change (Winger et al. 2019). Such changes, however, only confer a fitness benefit if animals arrive in the correct place at the correct time, with benefits only felt when local conditions match those required by the migrant. Rapid changes to the environment on the global scale might, therefore, be predicted to cause equivalently pronounced changes to the migratory routes of animals – though this would only be true if the mechanisms regulating migratory behaviour were either a) very plastic; b) transmitted socially; or c) capable of evolving rapidly. Since environmental change in the Anthropocene is particularly severe, understanding how migratory animals adapt (or fail to adapt) to these rapid changes is of both fundamental and practical importance (Conklin et al., 2021; Linssen et al., 2023; Lewin et al., 2024).

Perhaps one of the best-studied organisms in terms of continental-scale organisation of migratory strategy in both an evolutionary and behavioural context is the Eurasian blackcap (*Sylvia atricapilla*; ‘blackcap’). Blackcaps are very common breeders across western Eurasia, from Spain and Ireland in the west to Russia in the east. They are typically considered to be organised into three distinct migratory directions in autumn: a south-western migration in western Europe, a south-eastern migration in eastern Europe, and a more recently evolved northern/north-western route to Britain that occurs patchily across the central European range (Kopiec and Ozarowska 2012; Plummer et al. 2015; Tengholm et al. 2018; Delmore et al. 2020). These migratory routes have been shown to be genetically heritable, as has been investigated via selection and cross-breeding studies (Helbig 1991; Berthold et al. 1992; Pulido et al. 2001); and *in situ* the south-west and south-east routes are thought to form a narrow migratory divide at around 14°E (Pérez-Tris et al. 2004; Delmore et al. 2020). Birds from the area spanning the migratory divide apparently exhibit an intermediate southern migratory route in autumn, with south-west migrating and south-east migrating blackcap populations showing very little genetic differentiation (Pérez-Tris et al. 2004; Delmore et al. 2020; Delmore et al. 2023).

In a large-scale collaborative effort, we sought to characterise continental-scale organisation of blackcap migratory behaviour by tracking the individual migratory routes of blackcaps across the central European distribution range with focal efforts across the migratory divide, as well as the newly emergent NW phenotype with wintering sites in the British Isles (Delmore et al., 2020). As part of this study we deployed geolocator devices – archival devices used to determine position based on the light levels encountered through the year (Lisovski et al. 2020) – on birds caught in 2022 at two sites in north-eastern Poland. These sites were chosen to be clearly east of the hypothesised migratory divide (at 14°E), so as to increase the number of south-eastern migration routes in the dataset. This was because such routes were scarcely represented even from locations where the south-east phenotype was supposedly commonplace (Delmore et al. 2020). However, the tracks gathered (see Results) differed substantially from our expectations, and hence we sought out new data that represent spatial organisation of migratory strategies *and* entail a temporal component that allows to track this spatial organisation back in time. With this we tested the *a priori* expectation that blackcap migration was changing. Specifically, we used almost a century of citizen science ringing data, asking a) whether the continental-scale organisation of blackcap migration has changed and b), if so, why.

## Methods

### Geolocator deployment and processing

We equipped 50 adult male blackcaps with Migrate Technology P50Z11-11-COOL light-level geolocators via leg-loop harnesses constructed using elasticated plastic string. These devices weighed < 1.2 g including harness, and were set to record the maximum light level every ten minutes via a light sensor mounted on a 11 mm stalk (such that the sensor protruded above the feathers). Devices were deployed at two sites: Orzysz, Mazury; n =25, 53.81°N, 21.87°E; and Bielsk Podlaski, Białowieża Forest, Bielsk; n = 25, 52.67°N, 23.44°E. Five of these 50 devices were retrieved. Deployments and retrievals were made using 6 m/9 m mist nets (mesh size 16 mm) and male blackcap song as well as blackcap alarm call playback.

Since blackcaps spend the majority of their time in the undergrowth, precise light-level geolocation is challenging. As such, we avoided complex geolocation techniques (e.g. Rakhimberdiev et al. 2017) and instead used a simple threshold model (for a review, see Lisovski et al. 2020) to ascertain the average non-breeding position. We selected the sunlight elevation angle and threshold values that minimised the displacement of the estimated breeding location and the known breeding location (i.e. we used the breeding period of June-July as a ground-truthing period), which led to the selection of a sunlight elevation angle of - 3° and a threshold value of 3. The non-breeding location for each bird was ascertained as the median longitude/latitude of position over the period of boreal winter (December, January and February).

### Geolocation deployment ethics statement

Permits for equipping birds with geolocators, capturing and recapturing individuals, and collecting blood samples were obtained from the Local Ethics Committee in Poznań (resolutions no. 62/2020 and 73/2021) and the General Directorate for Environmental Protection (permit no. DZP-WG.6401.110.2020.AS).

### Selecting ringing records for analysis

The results of our geolocator deployments were intriguing, but by no means conclusive in determining how and why blackcap migratory organisation has changed. This is because any conclusions based on this data would necessarily be post-hoc, and hence we sought to investigate changes in blackcap migration with a new dataset that was substantially larger and allowed us to characterise potential changes over a larger temporal scale. This we did using ringing data gathered across Eastern Europe, obtained from the EURING based on a query for all blackcaps ringed and subsequently recovered, and this dataset was subsetted to retain only birds caught as breeders in Eastern Europe and recovered in the non-breeding period elsewhere. This was so as the birds retained for analysis did not include birds moving via Eastern Europe *en route* to/from other locations, and were instead reflective of genuine migratory movements.

Breeding birds were isolated based on breeding phenology and phenotype, with birds marked as ‘breeding’ if they were caught within a core breeding window of 15^th^ May – 15^th^ July or if they displayed a breeding phenotype (had a cloacal protuberance, a brood patch or were themselves ringed as a pullus in the nest). This window was chosen based on previous analyses of when European migrants were likely breeding and extremely unlikely migrating based on phenotypic data gathered via continent-wide ringing activity (Wynn et al. 2024).

Once breeding birds had been selected, they were then subsetted to include only birds caught at longitudes greater than 14°E (the position of the migratory divide determined via clinal analysis in Delmore et al., 2020). Birds breeding at greater than 25°E were also removed, since the phenotypes of blackcaps in Western Asia are poorly known *a priori*.

Each bird’s ringing record was paired to its respective recovery, and a migratory distance and direction vector calculated. Recoveries that were made during the breeding season were removed, since they were very unlikely to represent migratory movements, whilst ringing records made on migration removed by removing recoveries made within 500 km of the breeding site. This, it was hoped, would retain only records that reflect birds at (or very near to) the terminus of their migration.

### Assessing changes in migratory direction

Birds were assigned to one of three migratory orientation strategies (*sensu* Helbig 1991); ‘south east’ (90°-160°), ‘south’ (160°-200°) and ‘south west’ (200°-270°). To test whether the frequency of the use of each flyway was changing over time, we used a binomial logistic generalised linear model with probability of each migratory direction as a response variable in three separate models. Year was included as a predictor, alongside a term designed to control for variation associated with ringing effort. This was because any perceived changes in the use of a given flyway could reflect ringing effort on that flyway and not focal changes in migratory behaviour. For example, if birds were split equally between the two flyways, and ringing effort were to decrease in the east and increase in the west, then this would look like the eastern phenotype decreasing in probability and the western phenotype increasing in probability.

This term was calculated in a three-stage process. First, for each bird the probability of being recovered east of its present position as an ‘east-west odds’ was calculated. This was defined as the number of birds ringed (in the non-breeding period) east of the focal bird within 2 years’ of when the focal bird was ringed as a proportion of the total number of birds ringed over the same period:

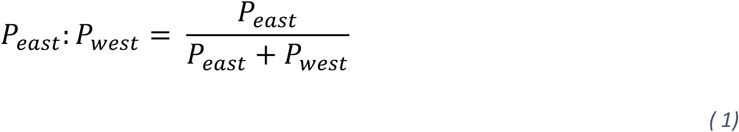

For example, for a bird ringed in 1963 at 52.67°N, 23.44°E, then the ‘east-west odds’ is the total number of birds ringed in the non-breeding period east of 23.44°E between 1961 and 1965 as a proportion of the total number of birds ringed in the non-breeding period between 1961 and 1965. If this figure was large (> 0.5), it would suggest eastern migratory routes were more likely to be detected. Conversely, if this figure were small (< 0.5) then this would suggest that western migratory routes were more likely to be detected. However, whilst this term is closely related to ringing effort, it is not solely influenced by ringing effort. Birds in the east must have a smaller east-west ratio, since there are fewer wintering sites east of them than there are to the west. Similarly, birds in the west must have a larger east-west ratio as there are fewer possible wintering sites to the west. To account for this, we then calculated for each bird an ‘east-west odds ratio’. First, we calculated for each bird an ‘expected east-west ratio’, which was defined as the median east-west ratio for each 5-longitudinal degree bin. Both the ‘east-west ratio’ and the ‘expected east-west ratio’ were converted to odds, and an odds ratio was calculated. This simplifies to:

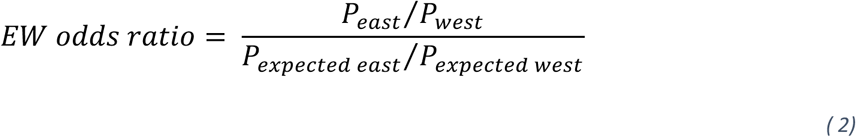

This ‘east-west odds ratio’, then, in essence captures differences in the probability of going east or west beyond those predicted simply by geographic position. This, in turn, is intrinsically linked to changes in detection probabilities associated with different migratory directions (caused primarily by differences in ringing effort). Because odds ratios distribute highly non-normally, the final stage of the process was to log-transform the resultant odds ratio. This creates a normally-distributed variable centred around zero – the ‘log east-west odds ratio’ – which was then included in the model, with the hope that it would account for variation associated with changes in ringing effort.

### Assessing changes in migratory duration/destination

We also sought to characterise differences between orientation strategies through time. Chiefly, we wanted to ascertain whether a) blackcaps were changing their wintering latitude through time, and b) whether different migratory strategies were altering their wintering latitudes to differing degrees. We tested this using linear regressions, and tested for significance with an ANOVA. To test whether birds following different orientation strategies were changing their wintering latitude per year, we regressed wintering latitude against year interacting with migratory strategy (east, west or south). A significant effect of year would suggest that birds were changing their latitude per year, whilst a significant effect of the interaction would suggest that the extent of this change differed between strategies. We repeated this analysis with migratory distance as the response variable, where again a significant effect of year would suggest that birds were changing their migratory distance whilst a significant interaction with strategy would suggest that different strategies were altering their migration in different ways.

As above, it is essential to include a term aimed at accounting for variance associated with ringing effort in the model, since changes in effort could confound with the variable of interest. For example, if the latitude of maximum ringing effort in the west of the range were to increase, this would cause the perceived wintering latitude of west-migrating to increase even if there were no change in the underlying migratory strategy. To do this, we calculated for each bird an ‘expected migratory distance’ and an ‘expected recovery latitude’ based on ringing effort. To do this we took each bird’s observed migratory bearing, and isolated all non-breeding ringing events that occurred within 250 km of the trajectory within 2 years of the focal bird’s ringing date. We then calculated the distance to each point and the latitude of each point, the median of which was determined to be the ‘expected migratory distance’ and an ‘expected recovery latitude’, respectively (see Figure S1). For example, if we again consider a bird for ringed in 1963 at 52.67°N, 23.44°E with a migratory direction of 105°, we can extrapolate that bearing and isolate the points within 250 km of a path along this bearing. These points can then be used to calculate an expected migratory distance/recovery latitude based on the distribution of ringing effort.

### Investigating the role of temperature in changing migratory behaviour

When testing for the environmental basis of any ecological change, it is tempting to ‘dredge’ parameter-space and test several environmental variables in the hope of finding the most likely causal relationship. However, we believe that given the complexity of the system no single environmental variable is likely to be directly and singly causal. For example, we might find that frost in December is the best predictor of changes in blackcap wintering behaviour. However, whether this means frost drives blackcap distribution or frost drives other factors that directly drive blackcap distribution – or both – is unclear; the analysis is still completely correlative, and dredging the dataset hasn’t changed that. As such, instead of testing multiple variables in the hope of determining a definitive, causative relationship, we instead sought to test a simple and singular hypothesis: that the observed temporal changes in migratory strategy over the last 100 years might be – directly or indirectly – correlated to changes in the winter temperature of Europe.

To test whether any changes in migratory behaviour reflected changes in temperature, we conducted two analyses aimed at assess a) whether temperature was changing in the blackcap wintering range (and whether the rate of change differed across longitudes), and b) whether changes in site occupancy were consistent with birds exclusively selecting traditionally ‘cold’ sites in years where they were warm. For both analyses, temperature data were sourced from the European Medium-range Weather Forecast ERA5 re-analysis dataset (downloaded from the Copernicus data centre; https://cds.climate.copernicus.eu/).

To address our first question, we regressed winter temperature – the mean temperature recorded over the months December to February – against, year, longitude and latitude for all locations at which blackcaps had been captured in the months December to February. The regression line was fitted via least means squares regression; significance was assessed with ANOVA; and the complete results table is available in the electronic supplementary materials.

To test whether birds were occupying traditionally ‘cold’ sites only in years where they were warm, we conducted a randomisation. If our hypothesis were true, we reasoned that the mean temperature encountered by blackcaps across their range would be warmer in the observed years than if the year of occupation were switched to a realistic but random alternative. This we did by sampling, with replacement, random years from the dataset and assigning them at random to different sites. For each iteration of this process, we calculated a mean temperature, which represented a realistic but random temperature value if the year a site was occupied was not influenced by temperature at that site. By repeating this process 1000 times, and computing the number of times the randomisation generated a temperature warmer than was observed, we could therefore calculate a p-value. This p-value was multiplied by two to account for the two-tailed nature of the randomisation.

## Results

### Geolocator analysis

Of the five birds re-caught the following year, four migrated west and one migrated south/south-east in autumn (see Figure 1). This was almost entirely at odds with the classical understanding of the geographic organisation of blackcap migration, which predicts that Eastern European breeders should migrate east (Delmore et al. 2020; Helbig 1991; Pérez-Tris et al. 2004).

**Figure 1:**
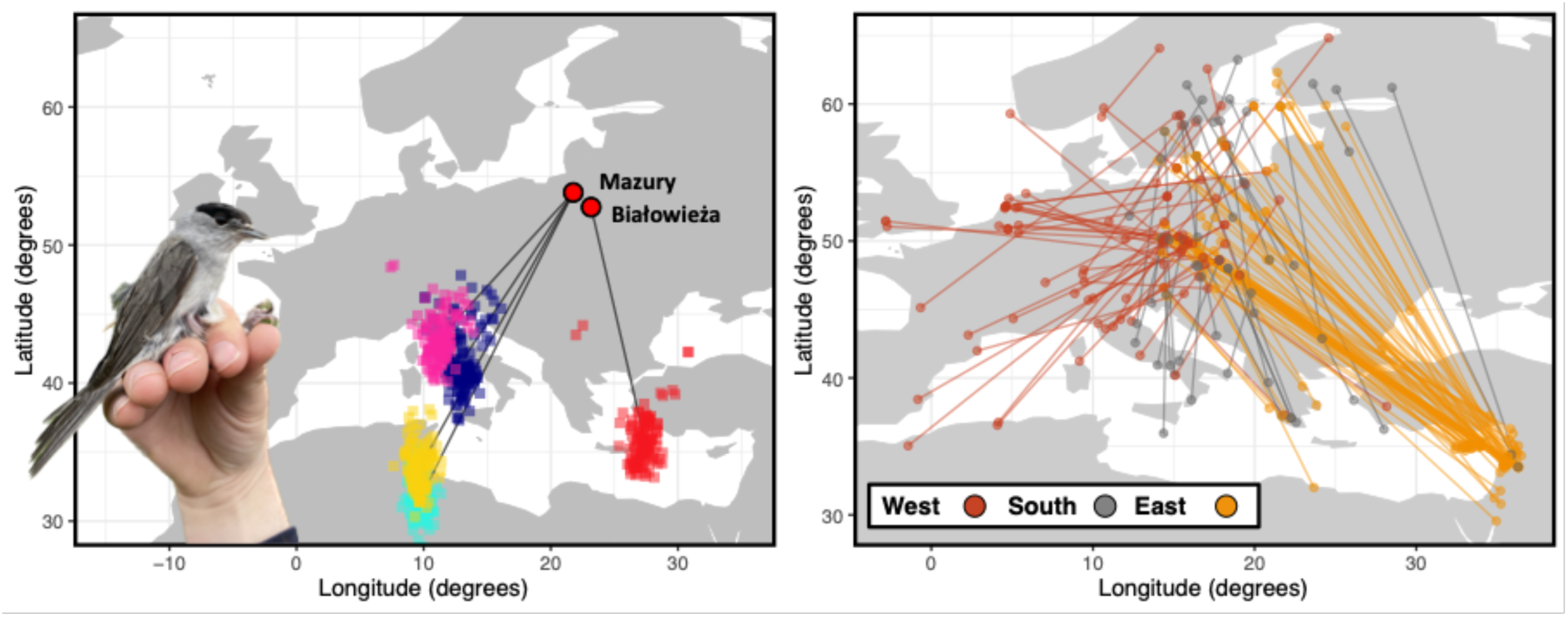
European blackcap migratory trajectories from eastern breeding sites. (left) the migratory trajectories of blackcap tracked using light-level geolocators from Białowieża (n = 1) and Mazury (n = 4) in north-eastern Poland in 2022/23. Points at the non-breeding site are coloured by individual, and represent each positional estimate in the months December, January and February. Lines join the non-breeding longitude/latitude to the known capture location. Inset photo shows a migratory blackcap carrying a recovered geolocator device. (right) all ringing recoveries of birds ringed in the breeding season east of the *a priori* estimated blackcap migratory divide (14° E); lines between ringing and recovery location are coloured by autumn migratory direction (west = red, south = grey and east = yellow).

### Ringing analysis

The extreme deviation between the migratory directions observed with geolocators and those expected based on previous studies is strongly indicative of Eastern European blackcaps changing migratory direction in the last century. However, such conclusions are necessarily post hoc, and hence we sought to use these data as a prior and instead conduct a more rigorous analysis using historically-gathered ringing data covering a large spatial and temporal scale.

First, we asked whether the probability of executing an eastern migratory route was reducing in Eastern Europe over time. Whilst we found no evidence for changing ringing effort driving changes in the detection probability of eastern migratory routes (binomial generalised linear regression; z = −0.674, p = 0.500; see Table 1), we found that the probability of using the eastern flyway had reduced markedly over the last century (binomial generalised linear regression; z = −4.05, p < 0.0001; Table 1). A year-on-year reduction of −0.0433 (± 0.0107) in the odds ratio suggested that eastern flyway utility had dropped from being near-fixed in the population in the early 20^th^ century to being almost unutilised in the 21^st^ century (see Figure 2), with this reduction in using of the eastern flyway matched by an increased use of the western flyway (binomial generalised linear regression; z = −5.02, p < 0.0001; see Table 1) but not the southern flyway (binomial generalised linear regression; z = 0.974, p = 0.330; see Table 1). Again, changes in ringing effort were implicated in neither change (western flyway; binomial generalised linear regression; z = 0.100, p = 0.921; southern flyway; binomial generalised linear regression; z = 0.424, p = 0.671), suggesting that overall longitudinal changes in ringing effort over the study period did not influence detection probabilities.

**Figure 2:**
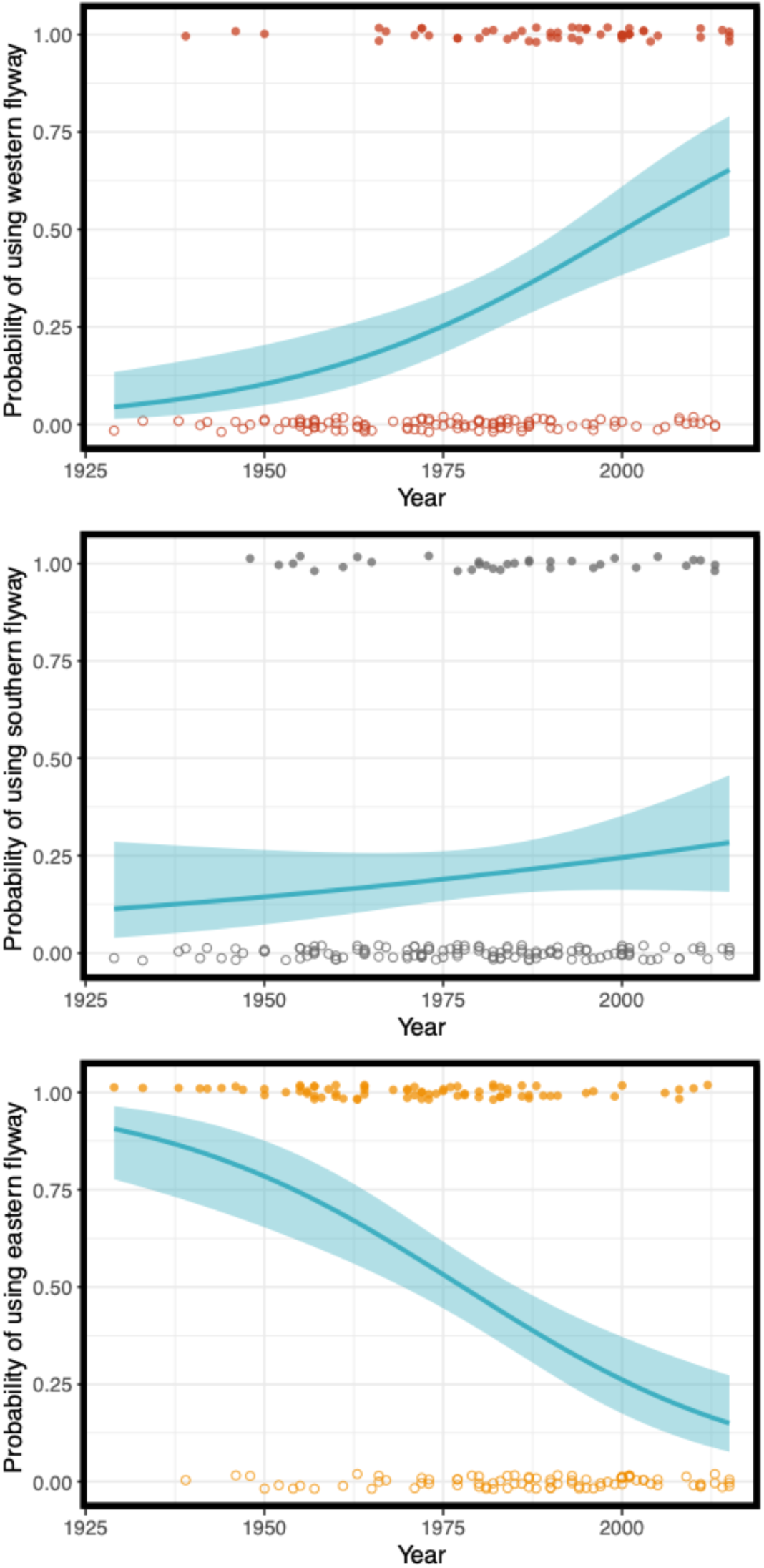
The declining usage of the eastern flyway in blackcaps over the last 100 years. Panels showing changes in the binomial probability of blackcaps migrating west (top), south (centre) or east (bottom) from blackcaps breeding at locations east of 14° E in autumn. Regression line shows predictions from a generalised linear model with a binomial distribution, with 95% confidence intervals shown about the regression line.

**Table 1:** Changes in the probability of blackcaps using different migratory strategies through time and with temperature. Generalised linear models detailing the changes in the use of different migratory strategies through time.

| Model | Term | Effect size (odds ratio) | Confidence interval (odds ratio) | Z-value | p-value |
| --- | --- | --- | --- | --- | --- |
| **Probability of migrating east ~ year + east-west odds ratio** | Intercept | 2.28 | 0.518 | **4.403** | **< 0.0001**<br>*** |
|  | Year | -0.0433 | 0.0107 | **-4.047** | **< 0.0001**<br>*** |
|  | East-west odds ratio | -0.180 | 0.267 | -0.674 | 0.500 |
| **Probability of migrating west ~ year + east-west odds ratio** | Intercept | -3.0565 | 0.609 | **-5.018** | **< 0.0001**<br>*** |
|  | Year | 0.0424 | 0.0116 | **3.641** | **< 0.0001**<br>*** |
|  | East-west odds ratio | 0.0243 | 0.244 | 0.100 | 0.921 |
| **Probability of migrating south ~ year + east-west odds ratio** | Intercept | -2.065 | 0.579 | **-3.569** | **<0.001**<br>*** |
|  | Year | 0.011 | 0.0113 | 0.974 | 0.330 |
|  | East-west odds ratio | 0.112 | 0.263 | 0.424 | 0.671 |

It is possible that the shift from eastwards to westwards migration trajectories in autumn could reflect anthropogenic climate change allowing for the possibility of wintering at more northerly latitudes in the west. Such opportunities would not necessarily manifest in the east, owing to the Adriatic Sea and Carpathian Mountains occupying the equivalent latitudes, making a western migratory route potentially selectively advantageous. To test this, we regressed latitude and migratory distance in separate models against year interacting with migratory strategy (west, east or south) in a linear regression. Again, we included a term designed to account for variation in the detected migratory latitude/distance owing to ringing effort (see Methods). When considering migratory distance, we found that migratory distance as perceived using ringing data was significantly predicted the distribution of ringing effort (F = 60.7, p < 0.0001; see Table 2; see Methods) but that, when these changes in ringing effort were accounted for, the distance between ringing and recovery was significantly predicted by migratory strategy (F = 99.3, p < 0.0001; see Table 2), year (F = 115.2, p < 0.0001; see Table 2) and the interaction between strategy and year (F = 4.92, p < 0.01; see Table 2; see Figure 3). This suggests that migratory distance varied between migratory strategies, and that changes in migratory distance over time differed between strategies. We found that the recovery latitude of migratory birds mirrored this result perfectly, with recovery latitude significantly predicted by the latitudinal distribution of ringing effort along each flyway (F = 72.8, p < 0.0001; see Table 2; see Methods), migratory strategy (F = 136, p < 0.0001; see Table 2), year (F = 135.677, p < 0.0001; see Table 2) and the interaction between strategy and year (F = 4.81, p < 0.01; see Table 2; see Figure 3). Post-hoc comparison suggested that eastwards migratory routes were not reducing significantly year-on-year (0.834 km/year ± 2.837 km/year), in contrast to the significant and rapid route attenuation seen in west-migrating (−8.30 km/year ± 4.62 km/year) and south-migrating populations (−15.3 km/year ± 5.04 km/year). Correspondingly, post-hoc analysis revealed a non-significant change in recovery latitude per year in east-migrating blackcaps (−0.0180° ± 0.0250°), whilst wintering latitude increased by 0.0891° (± 0.041°) a year in west-migrating birds and 0.125° (± 0.0450°) per year in south-migrating birds. This implies, therefore, a) that migratory routes to the west and south are shorter, and b) that in contrast with eastern migrants, these routes have gotten shorter year-on-year over the past century.

**Figure 3:**
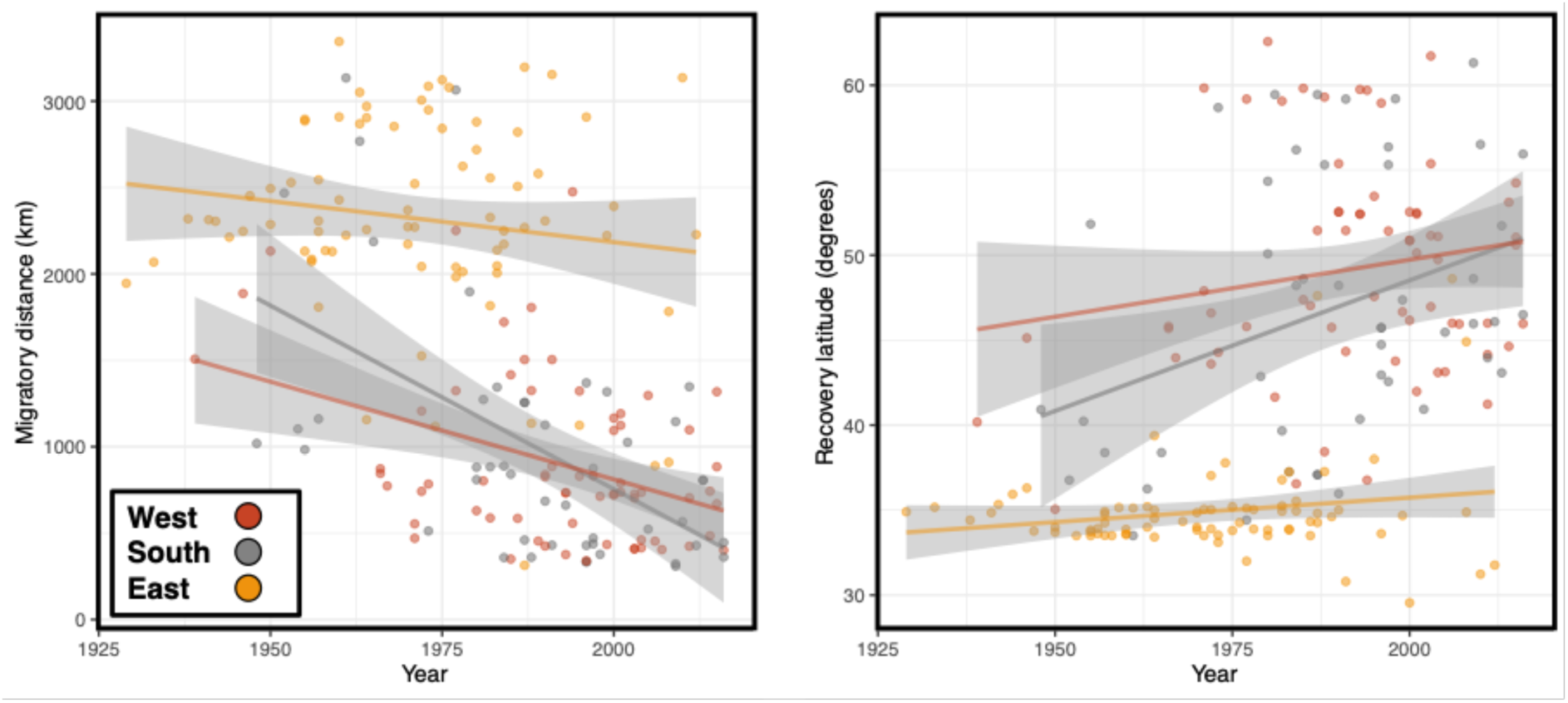
changes in migratory distance and destination through time for different migratory strategies. (left) The migratory distances of blackcap migration since 1925 coloured by migratory autumn direction (west = red, south = grey and east = yellow). (right) The changes in latitude at which blackcaps breeding east of 14°E are recovered since 1925 coloured by autumn migratory direction.

**Table 2:** Changes in migratory distance and recovery latitude for different blackcap migratory strategies. Linear models detailing the relationship between migratory strategy, year and migratory distance/recovery latitude. Linear models were assessed for significance using ANOVA statistics, and hence effect sizes and confidence intervals were not used in the estimates of statistical significance presented here. To avoid confusion, effect sizes and confidence intervals are presented in the main text.

| Model | Term | F-value | p-value |
| --- | --- | --- | --- |
| **Migratory distance ~ year \* migratory strategy + expected migratory distance** | Migratory strategy | 99.319 | **< 0.0001**<br>*** |
|  | Year | 115.253 | **< 0.0001**<br>*** |
|  | Year * Migratory strategy | 4.923 | **< 0.01**<br>** |
|  | Expected migratory distance | 72.841 | **< 0.0001**<br>*** |
| **Recovery latitude ~ year \* migratory strategy + expected recovery latitude** | Migratory strategy | 136.610 | **< 0.0001**<br>*** |
|  | Year | 135.678 | **< 0.0001**<br>*** |
|  | Year * Migratory strategy | 4.811 | **< 0.01**<br>** |
|  | Expected recovery latitude | 60.712 | **< 0.0001**<br>*** |

Given the extremely rapid changes observed using ringing data, we reasoned that it was possible that such changes are explained directly by anthropogenic climate change. As such, we performed two analyses. As mentioned in the methods, we chose not to ‘dredge’ environmental variables and instead perform a single, hypothesis test on the most interesting candidate; winter temperature change. First, we investigated whether winter temperature has changed in the blackcap wintering range and if so, whether this changed was more pronounced in the west than the east (consistent with birds switching routes to utilise warmer conditions at higher latitude in the west). We found that that temperature was increasing significantly across the blackcap range (F = 1835, p < 0.001; see Table S1), with temperature changes at the mean blackcap wintering longitude and latitude changing by 0.0684° C ([95% CI] ± 0.0435° C) per year. The interaction with longitude was also highly significant (F = 131, p < 0.001; see Table S1), with the effect of year reduced by −0.0135° C (±0.000484° C) for each further degree east moved. This suggests that temperature is increasing at a greater rate in the west of the blackcap range, consistent with the east-to-west shift being caused at least in part by changes in temperature.

Second, we tested whether the temperatures in at blackcap wintering sites were warmer in the year that the site was occupied than would be expected by chance. If this were true, we reasoned, this would be supportive of birds selecting traditionally ‘cold’ sites only in years where they were warmer. To test this, we calculated the mean temperature recorded across all blackcap wintering sites in the years the sites were active. We then randomised the year-site pairings – to simulate site use in other years – 1000 times to create a realistic but random expectation of what temperature would be if birds selected the same sites but in different years (see Methods). When we did this, we found that the temperature birds experienced was significantly warmer than if they’d used the same sites in randomly selected years (p < 0.001), consistent with the hypothesis that birds selected colder wintering sites in years that were warmer.

## Discussion

Our results suggest that blackcaps breeding to the east of the previously clearly characterised location of the divide have almost completely shifted in their migratory orientation over the past century. Instead of following their traditional route to the south-east (Helbig 1991), ringing data confirm our geolocator data and suggest that almost all birds now use a shorter-distance route to the south-west. This route appears to facilitate birds wintering at higher latitudes, with these changes correlated with changes in the temperature over the wintering period. Whilst, as with any correlative analysis, it is possible that the correlation with temperature is not causative. Indeed, given the myriad possible impacts of temperature we might suggest it is perhaps even more likely that the effect of temperature is indirect (Siddiqi-Davies et al. 2024). However, given that seasonality-induced changes in temperature are seen as a primary driver of seasonal migration (Winger et al. 2019), we believe that the effect of temperature is unlikely caused by a confounding variable, and hence is of interest when considering the evolution of migration.

When reflecting on rapid changes in migratory behaviour, it is essential that we understand the mechanisms via which animals derive migratory information. In many animals, migratory movements are underpinned by information learnt either via asocial learning (often thought to be associative ‘trial and error learning’; Thorup et al. 2003; Campioni et al. 2020; Wynn et al. 2020) or through cultural transmission (social learning from experienced conspecifics; Mueller et al. 2013; Abrahms et al. 2021; Byholm et al. 2022). Migrants that make extensive use of learnt information, primarily long-lived species, have been shown to be capable of flexibly altering their migratory routes in response to a changing climate (Conklin et al. 2021; Linssen et al. 2023; Lewin et al. 2024), though this mechanism seems unlikely to explain changes in blackcap migration. This is because blackcaps are solitary migrants with short generation times, where migratory information is thought to be inherited genetically.

Extensive evidence for the genetic inheritance of songbird migration comes from both lab-based assays and selection and cross-breeding experiments – involving both blackcaps and related taxa – (Berthold and Querner 1981; Helbig 1991; Pulido et al. 2001; Zolotareva et al. 2021; Wynn et al. 2023), as well as *in situ* field observations and manipulations (Perdeck 1958; Harris 1970; Chernetsov et al. 2004; Thorup et al. 2007; Handel and Gill 2010; Yoda et al. 2017; Pot et al. 2024). Whilst recent studies suggest that a surprising number of such species in the Americas are vocal at night (Van Doren et al. 2025), it is unclear what role such vocalisations play in informing orientation. Indeed, it is unclear how individuals might differentiate informed and uninformed individuals and hence make decisions about what information to integrate, and there is currently no evidence that blackcaps make such vocalisations. As such, given the weight of evidence that points to a genetic inheritance of the migratory trajectory, we might suggest that the changes observed here are most likely to reflect microevolutionary changes in the genetically inherited migratory information used by blackcaps and – given the speed of these changes – we might assert that this is likely to reflect changes in existing allele frequency rather than any kind of *de novo* mutation.

These changes appear to facilitate shorter distance migration owing to increasing winter temperatures at higher latitudes, suggesting that as the climate changes the same continental formation can might give rise to different selection pressures in different climatic conditions (e.g. Voelker and Light 2011). This, in turn, appears to cause changes in the continental-scale organisation of migratory movements, such as the changes in blackcap migration observed here. Taken together, this suggests that continental-scale migratory organisation can, at least in some species, evolve surprisingly rapidly in response to anthropogenic changes.

Changes in migratory behaviour mediated via behavioural plasticity has been observed in several long-lived taxa (Conklin et al. 2021; Linssen et al. 2023; Lewin et al. 2024). Whilst this has been postulated as a potential mechanism via which long-lived migratory animals might buffer again anthropogenic environmental alteration, the results discussed above suggest that extremely large populations of birds with short generation times and heavy reliance on genetically inherited material might also respond rapidly to a changing environment. Given the speed of the response reported here for blackcaps, we might suggest that rapid evolution likely makes use of standing genetic variation in driving changes in the migratory phenotype. As with any correlative study using data gathered from free-flying birds, further verification either from experimental paradigms (e.g. *in situ* cross breeding) or from multiple other independent data sources is essential. Nonetheless, our study highlights the importance of large-scale longitudinal data collection in the study of how animal behaviour can change on a continental scale through time, and highlights the importance of understanding migratory behaviour such that we can understand the potential impacts of large-scale environmental changes.

## Supporting information

Supplemental PDF containing peripheral results and figures.

## Acknowledgments

We are grateful to the European Union for Bird Ringing (EURING) that made the recovery data available through the EURING databank and to the many ringers and ringing scheme staff, who have gathered and prepared the data. Funding was gratefully received through the Max Planck Society (MPRG grant MFFALIMN0001 to ML), the DFG (project Nav05 within SFB 1372 – Magnetoreception and Navigation in Vertebrates, project no 395940726, to ML), and the Ministry for Science and Culture of Lower Saxony. Permits for equipping birds with geolocators, capturing and recapturing individuals, and collecting blood samples were obtained from the Local Ethics Committee in Poznań (resolutions no. 62/2020 and 73/2021) and the General Directorate for Environmental Protection (permit no. DZP-WG.6401.110.2020.AS).

## Notes

https://zenodo.org/records/18674081?token=eyJhbGciOiJIUzUxMiJ9.eyJpZCI6ImU3MjZiYzA0LTgyMzItNDcxZC1hMDY4LTFjN2QwYzM5YzY5NCIsImRhdGEiOnt9LCJyYW5kb20iOiIxYmViMzMzN2ViODA3YzYyNmI3OTVhNTQxYjgyMmVjMCJ9.I8FsCUXnTMXuxXWoeS6bitOS-eJQ0PuzhvZIT3hs4DItl5vGDl6JFlY03oyRSm08HGgrHOO5c1yzbuTxZLcfcA

### Competing Interest Statement

The authors have declared no competing interest.

