## Supplemental PDF containing peripheral results and figures. for "Rapid and divergent changes in the continental-scale organisation of a short-lived songbird’s migratory strategy"

*Electronic supplementary material*

Joe Wynn[1,2], Monika Broniszewska[3], Alice Edney[4], Tania Garrido-Garduño[5], Joe Morford[6], Michał Polakowski[3], Robert Rollins[2], Pablo Salmon[2], Oscar Vedder[2] and Miriam Liedvogel[2,7,8]

| | *Estimate* | *Std. Error* | *Confidence interval* | *F value* | *Pr(>F)* |
|---|---|---|---|---|---|
| ***Intercept*** | 9.699525301 | 0.021944452 | 0.043450015 | NA | NA |
| ***Year*** | 0.068384322 | 0.002223705 | 0.004402936 | 1835.880562 | < 0.0001 *** |
| ***Mean-centred longitude*** | -0.124139054 | 0.003755763 | 0.00743641 | 5735.210761 | < 0.0001 *** |
| ***Mean-centred latitude*** | -0.879541689 | 0.005481068 | 0.010852515 | 29505.93583 | < 0.0001 *** |
| ***Mean-centred year * Mean-centred longitude*** | -0.003516827 | 0.000299101 | 0.000592219 | 131.3807557 | < 0.0001 *** |
| ***Mean-centred year * Mean-centred latitude*** | -0.013556695 | 0.000484483 | 0.000959277 | 1143.548185 | < 0.0001 *** |
| ***Mean-centred longitude * Mean-centred latitude*** | -0.008889071 | 0.000721986 | 0.001429533 | 197.4304039 | < 0.0001 *** |
| ***Mean-centred year * Mean-centred longitude * Mean-centred latitude*** | 0.000412556 | 6.02156E-05 | 0.000119227 | 46.94046953 | < 0.0001 *** |

**Table S1: Year-on-year changes in temperature across the blackcap wintering range.** The effect of year, longitude and latitude on wintering site temperature from 1940-2016 for all recorded blackcap wintering sites.

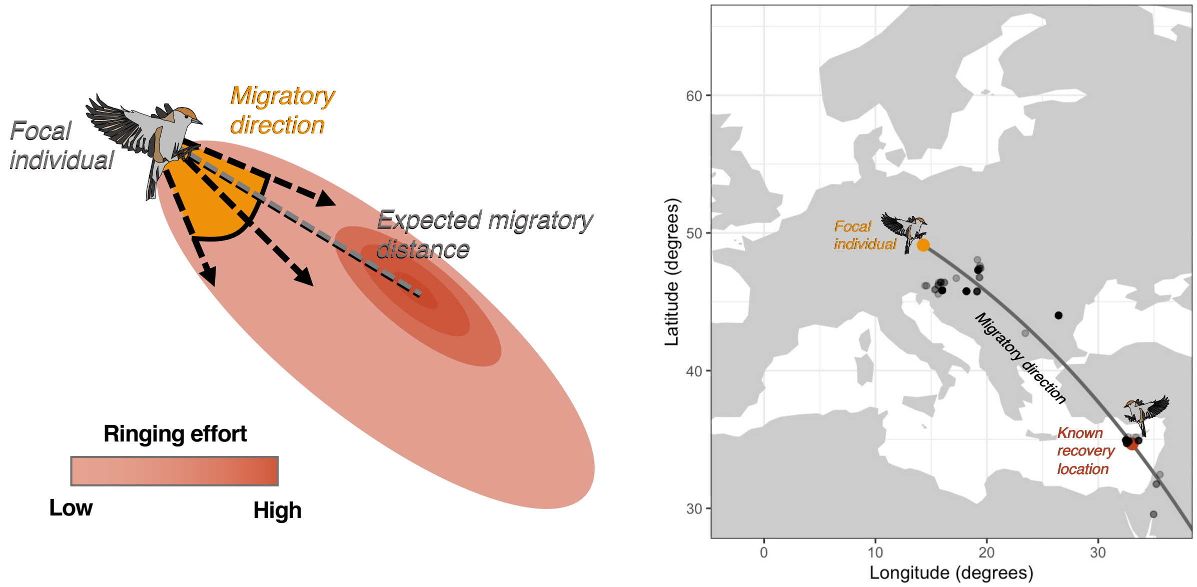


**Figure S1: Calculating the 'expected migratory distance' based on ringing effort.** (left) A schematic showing how, if all birds migrated the same distance, changes in ringing effort might bias estimates of migratory distance. (right) An exemplar case showing how the expected migratory distance can be calculated by examining ringing effort along the trajectory between where a focal individual was ringed and recovered.
